# Inhibition of *Klebsiella pneumonia* by a Novel Strain of *Paenibacillus*

**DOI:** 10.1101/482950

**Authors:** Muhan Yang, Huijuan Su, Xinru Cheng, Xiaobo Li, Huihui Lian, Yuqi Wang, Yang Wang

## Abstract

*Klebsiella pneumoniae* is the causative agent of Klebsiella pneumonia and enteritis, and the prevalence of antibiotic resistant strains is becoming a serious public health problem. In this study, we isolated a novel strain of *Paenibacillus polymyxa* from the fecal extracts of healthy dogs that were challenged with *K. pneumoniae*. By combination of transposon mutagenesis and metabolic analysis, a nonribosomal peptide synthase gene cluster was identified to be involved in the antagonism, and the molecular weight of the compound was 1168.38 g/mol. These findings will enlarge the arsenal against drug-resistant pathogens.

## Introduction

*Klebsiella pneumoniae* belongs to the Enterobacteriaceae(1) and is a conditional pathogen with worldwide distribution. It is a resident bacterium of animal respiratory and intestinal tracts and can infect not only animals but also humans(2, 3). In general, *K. pneumoniae* are notorious regarding nosocomial infection and immunodeficiency infection(4). The main symptoms of infection are bacteremia, liver abscesses, endophthalmitis, meningitis, and metastatic infections(5–11). Aminoglycosides and cephalosporins are commonly used to treat *K. pneumoniae* infection(12). However, in recent years, the abuse of antibiotics resulted in the emergence of drug-resistant *K. pneumoniae* strains carrying with KPC(*Klebsiella pneumonia* carbapanemase) gene, VIM(Verona integrin-encoded-metallo-β-lactamases) gene, IMP(Imipenem metallo-β-lactamase) gene and NDM-1(New Delhi metallo-β-lactamase 1) gene(13). Notably, the NDM-1 strains are resistant to vast majority of antibiotics, which include not only cephalosporins, but also the carbapenes, the last resort antibiotic for the treatment of multidrug-resistant Gram negative bacteria(14), thus imposing a huge threat to public health.

The preferred strategies for the combat against superbugs mainly relied on the discovery of novel antibiotics and strain-specific phages. Although phage therapy has attracted more and more attentions in nowadays, antibiotic therapy remains the staple treatment. Prokaryotic secondary metabolites are a rich source of antibiotics (e.g., erythromycin from *Saccharopolyspora erythraea*)(15). Screening for novel active compounds from microorganisms is further accelerated due to the advance in genome sequencing and genome mining(16).

This study was driven by a severe outbreak of *K. pneumoniae* enteritis in a kennel of American bully in Nanjing, which was insensitive to cephalosporins nor carbapenes. The survived dogs inspired us to isolate the beneficial bacteria that antagonized *K. pneumoniae.* Through a large work of screening, we successfully isolated the strain WY54, which showed potent inhibitory activity towards *Klebsiella pneumoniae*.

## Materials and Methods

### Pathogen isolation and identification

Symptoms of vomiting, diarrhea, and bloody stools broke out in a large kennel. Even after treatment with the third generation cephalosporins, those symptoms remained. Ultimately, those sick dogs died several days later. Small intestinal specimens of dead dogs were collected for bacteria analysis. Then aerobic cultures were plated on blood agar plate and incubated at 37℃, and the colonies were uniform and were all resistant to cephalosporins.

Eight colonies were picked up for PCR amplification using the primers targeting 16S rRNA (KP Forward and KP Reverse). The PCR products were gel purified and cloning into the pGEM-T Easy vector and sequenced with universal primer M13 forward and M13 reverse. Sequence analysis revealed that all strains were identical, and here designated as *Klebsiella pneumoniae* MH18 (Genbank Accession: MH698490). The putative gene(s) encoding the carbapenemase were PCR amplified using the degenerate primers listed in table 1.

**Table.1.**
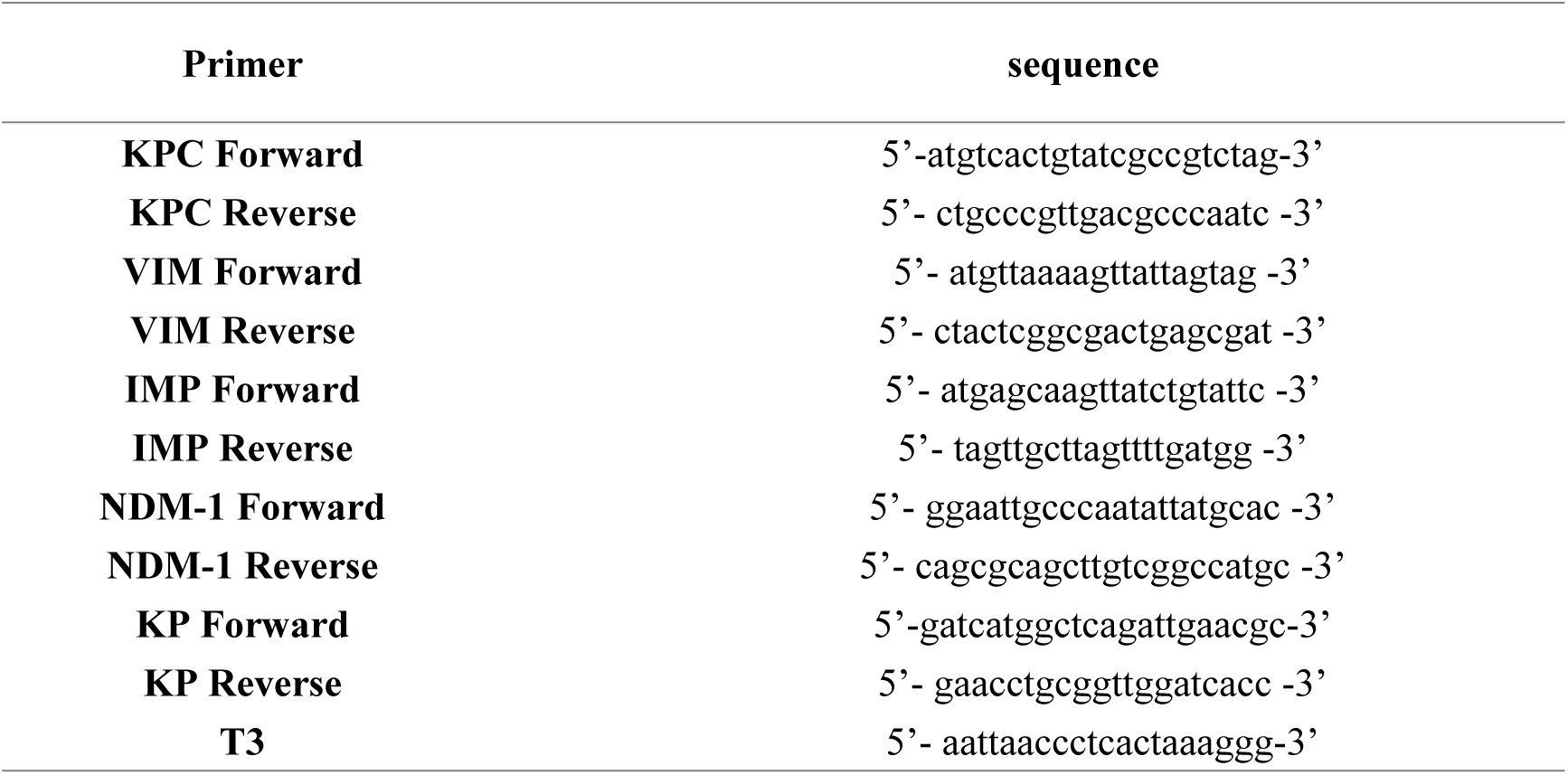
Primer sequences

### Screening for antagonistic bacteria against strain MH18

Bioassay against *Klebsiella pneumoniae* MH18 were carried out according to Yang(17), with candidate antagonistic strains isolated from the gut of healthy dogs in the disease-breaking kennel. After a large number of plate confrontation culture experiments, one strain showed strong inhibition activity against strain MH18. Biochemical analysis and 16S rRNA sequence determined that the antagonistic strain belongs to *Paenibacillus polymyxa*, and here designated as WY54 (Genbank Accession: MH698489).

### Transposon mutagenesis and gene cluster rescue

Transposon mutagenesis of *Paenibacillus polymyxa* WY54 was carried out as described by Dominique Missiakas(18), with minor modifications. The plasmids pFA545 and pBursa were sequentially transformed into strain WY54, and the resulting transformants were spread on LB containing 10 μg/ml of erythromycin and incubated overnight at 30 ℃. The single colonies which were isolated from LB_erm_ were suspended in 150 μl of deionized sterile water and spread on LB_erm_. The colonies were incubated at 43 ℃, lasting for two days, to select desired mutants.

Genomic DNA of the mutant strain WY54-MT7 that lost activity against *Klebsiella pneumoniae* MH18 was employed to construct a cosmid library using SuperCos1 (Invitrogen) as the vector(17), and the transformants were selected with erythromycin to rescue the fragments harboring transposon. Cosmids harvested were sequenced starting from primer T3, and the sequences were then assembled for antiSMASH analysis(19) (Genbank Accession:MH999500-MH999504), as well as A-domain analysis using the NRPSpredictor2(20).

### Isolation and identification of the antagonistic compound

*Paenibacillus polymyxa* WY54 wild type and mutant strain WY54-MT7 were cultured for high performance liquid chromatography (HPLC). After filtering through a 0.45 μm syringe filter, a 10 μl sample was injected onto a HyPURITY C18 column (5 μm, 150×4.6 mm) attached to HPLC. Separation was achieved using a 45 min linear gradient from 10% to 50% acetonitrile at a flow rate of 1.0 ml/min. The UV detection wavelength was 212 nm. The main peak that disappeared in the mutant strain was collected for bioassay against *Klebsiella pneumonia* MH18. Concentrated samples with bioactive activity obtained by HPLC were analyzed by mass spectrometry on the following condition: Ion source: ESI source; Capillary voltage: 4000 V; Dissociation voltage: 20 V; Infusion pump flow rate: 10 μl/min.

## Results

### Pathogen identification and biochemical analysis

Bacteria suspension made from the feces of sick dogs were plated on MacConkey agar, and all colonies were mucoid, rod-shaped and Gram-negative after staining. 16S rRNA sequencing confirmed their uniformity and demonstrated a 99% similarity with *Klebsiella pneumoniae* (designated as strain MH18). Antibiotic resistance test showed that strain MH18 was resistant to all the antibiotics tested, including tetracycline, apramycin, kanamycin, chloramphenicol, bleomycin, ampicillin, cephalosporin and meropenem (Fig 1). To detect the gene that confers resistance to carbapenem, the genomic DNA of strain MH18 was extracted for PCR amplification using the degenerate primers that target carbapenemase genes. Among the 4 amplicons, the primers targeting NDM-1 gene showed a strong positive band (Fig 2), and BLAST analysis indicated that the sequence shared a 99% similarity with the NDM-1 gene (Genbank Accession: NC_019153.1).

**Fig 1.**
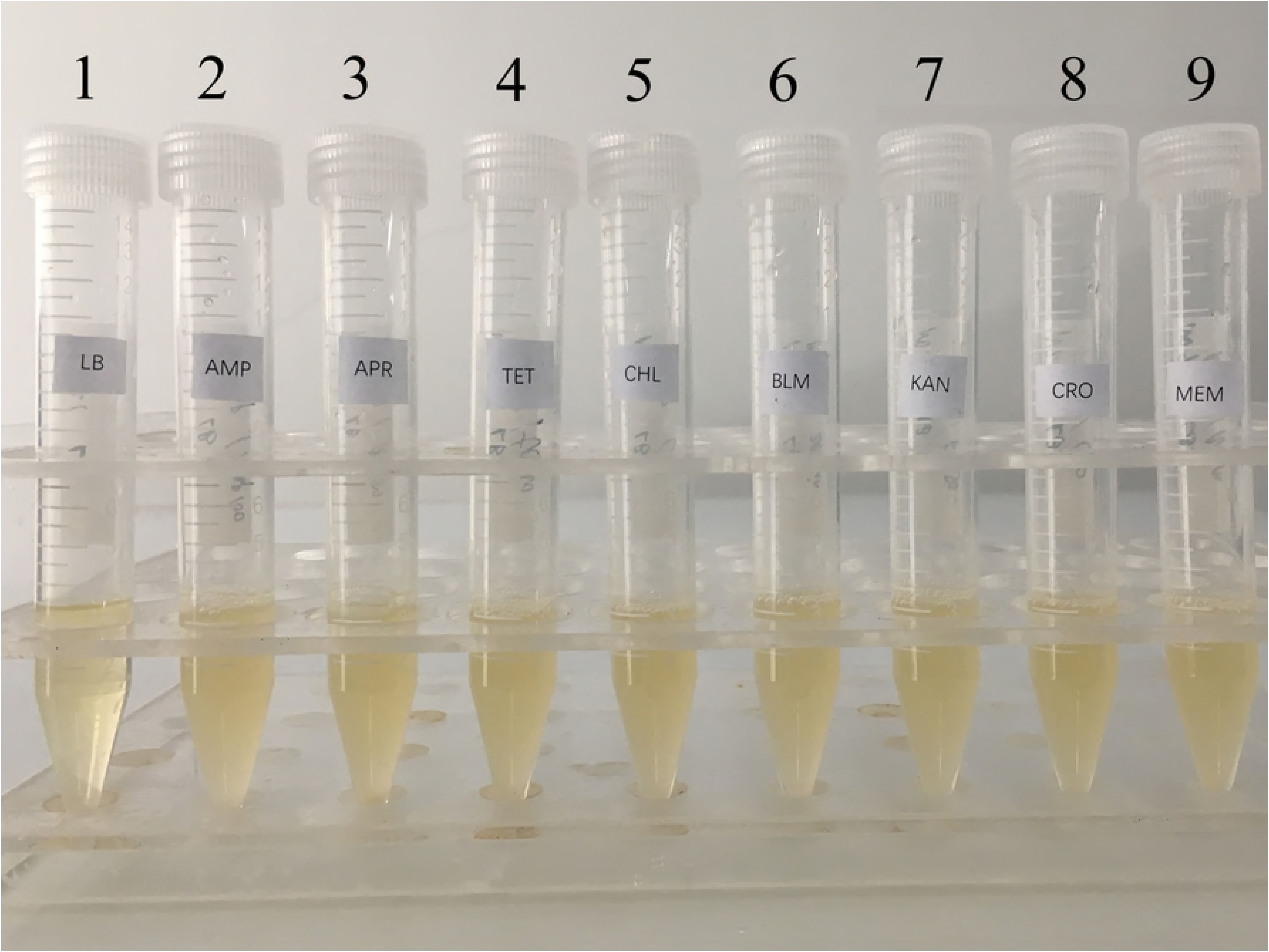
Antibiotic resistance test of *Klebsiella pneumonia* MH18 (1) CK (2) Ampicillin (3) Apramycin (4) Tetracycline (5) Chloramphenicol (6) Bleomycin (7) Kanamycin (8) Ceftriaxone Sodium (9) Meropenem

**Fig 2.**
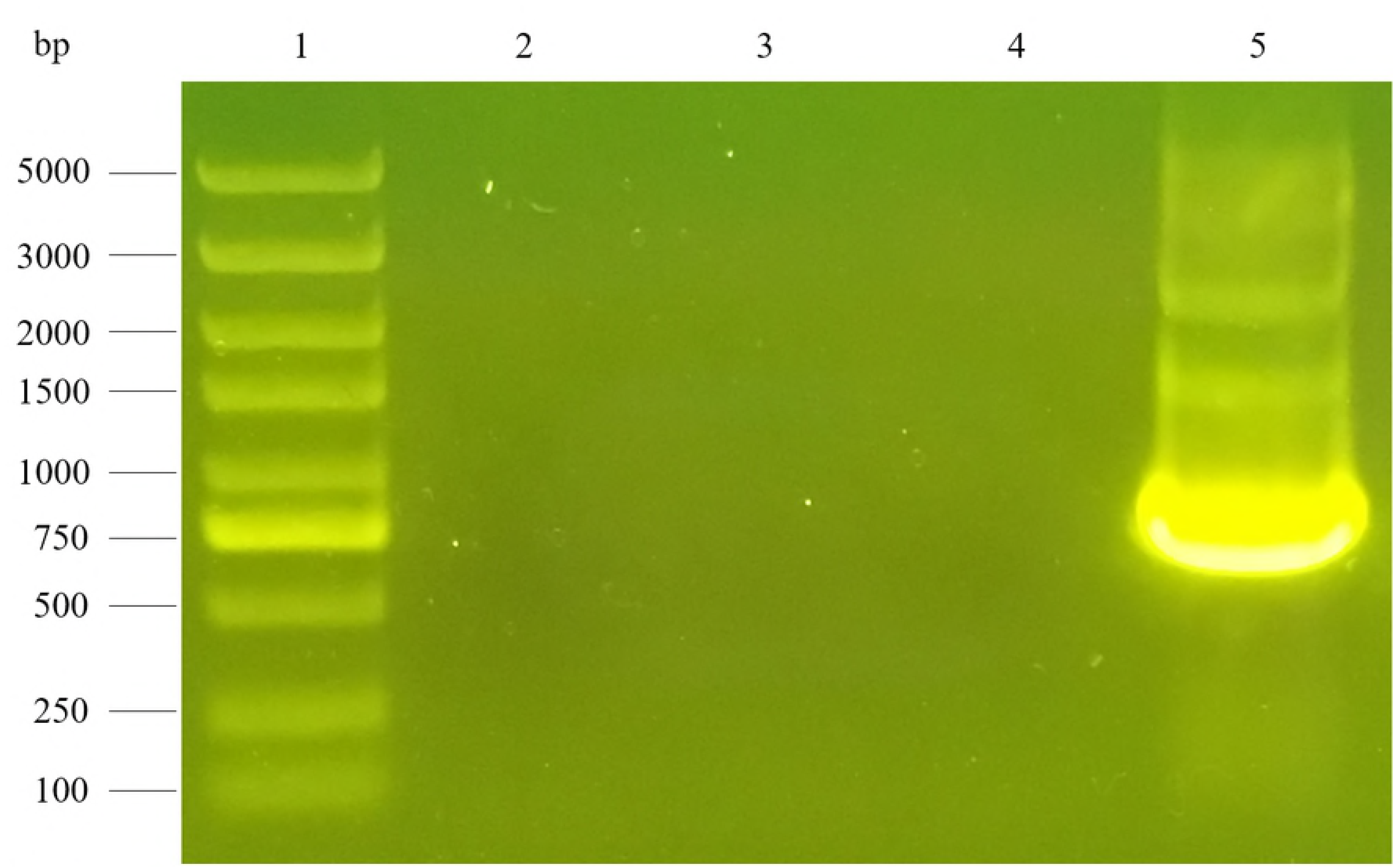
PCR amplification of the putative carbapenemase genes 1. KPC; 2. VIM; 3. IMP; 4. NDM-1. The DNA marker used is 1 kb DNA ladder.

### Antagonistic screening with *K. pneumoniae*

On contrast to *Bacillus subtilis* 168, strain *Paeniabcillus polymyxa* WY54 was antagonistic toward *K. pneumoniae* MH18. After 24 h of confrontation cultivation, an inhibition zone of about 9 mm in diameter was formed around strain WY54 on the lawn of *K. pneumoniae* MH18. The phylogenetic tree of strain WY54 was shown in Figure 3. The evolutionary history was inferred using the Neighbor-Joining method(21), and the evolutionary analyses were conducted in MEGA7(22). *Paeniabcillus polymyxa* are known for their ability to produce various kinds of secondary metabolites, which played important roles in the biological control of bacterial and fungal diseases(23).

**Fig 3.**
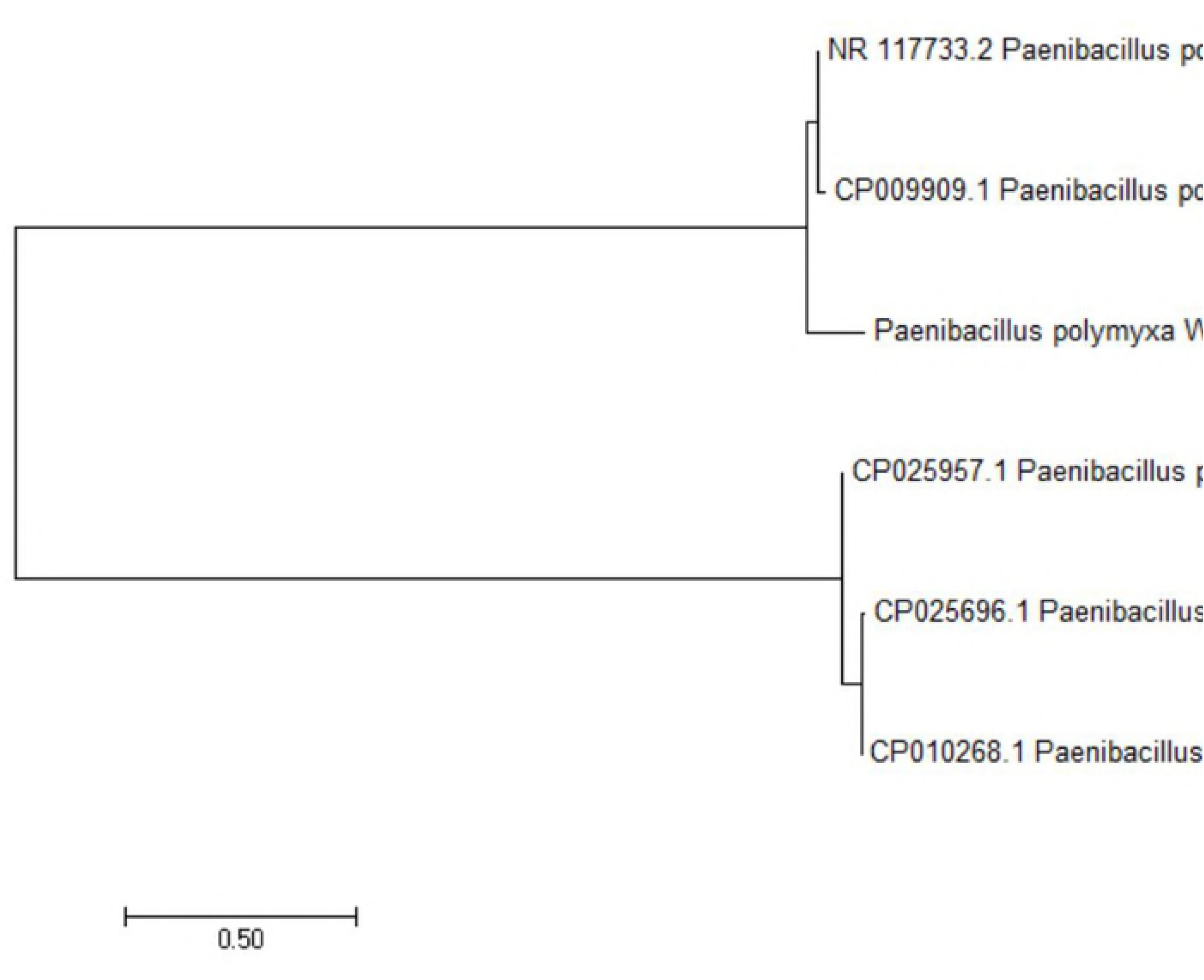
Evolutionary relationships of WY54

### Characterization of the gene cluster responsible for the antagonism

The wild type strain of *P. polymyxa* WY54 and its mutant were applied for bioassay against strain MH18 (Fig 4). Cosmid clones of the transposon mutant WY54-MT7 were sequenced and the fragments overlapping each other were aligned to obtain a gene cluster of ~40 kb in length. The gene cluster was analysed using BLAST, antiSMASH(19), and NRPSpredictor2(20). BLAST result suggested the genes encode non-ribosomal peptide synthases (NRPS), and the antiSMASH data predicted a depsipeptide with a 10-amino acid core structure (Fig 5.B). Further analysis with NRPSPredictor2 showed that the adenylation domain of module 1 recognized the rare amino acid Dab but not Ala, as shown in antiSMASH data (Fig 5.B), thus drawing the structure of the antagonistic compound closer to polymyxin E1(24).

**Fig 4.**
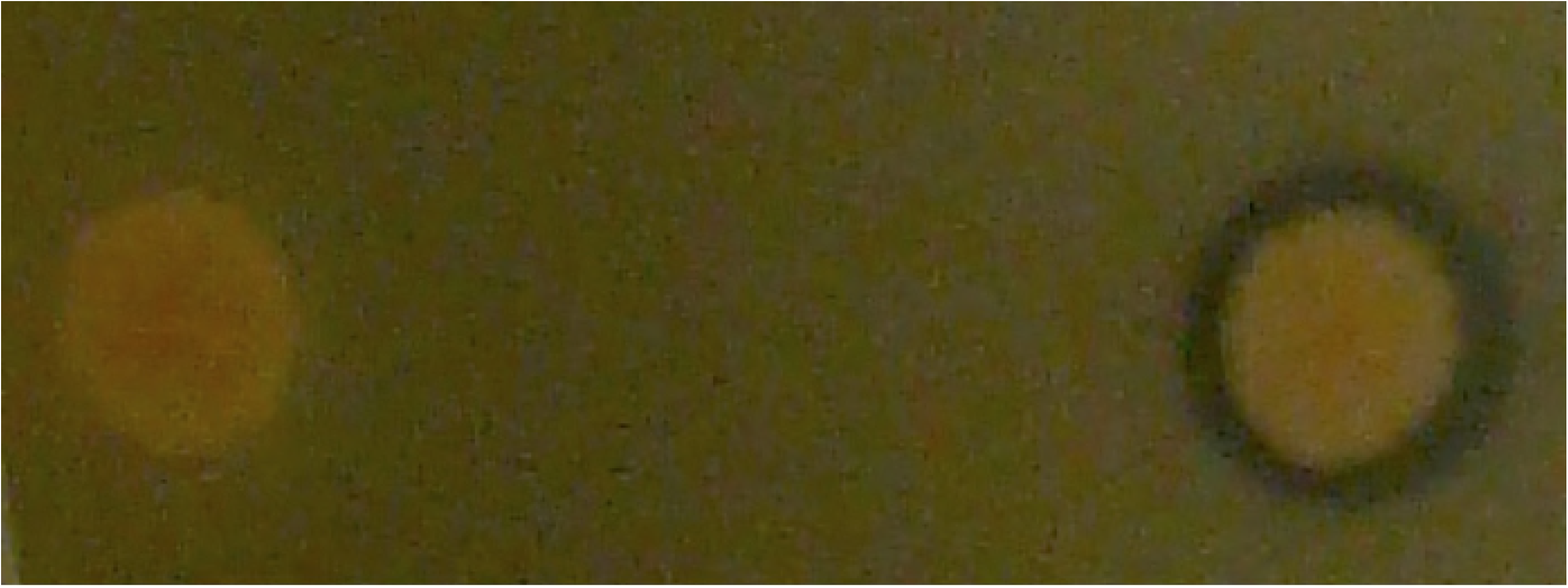
Bioassay of strain WY54-MT7 and strain WY54 against strain MH18

**Fig 5.**
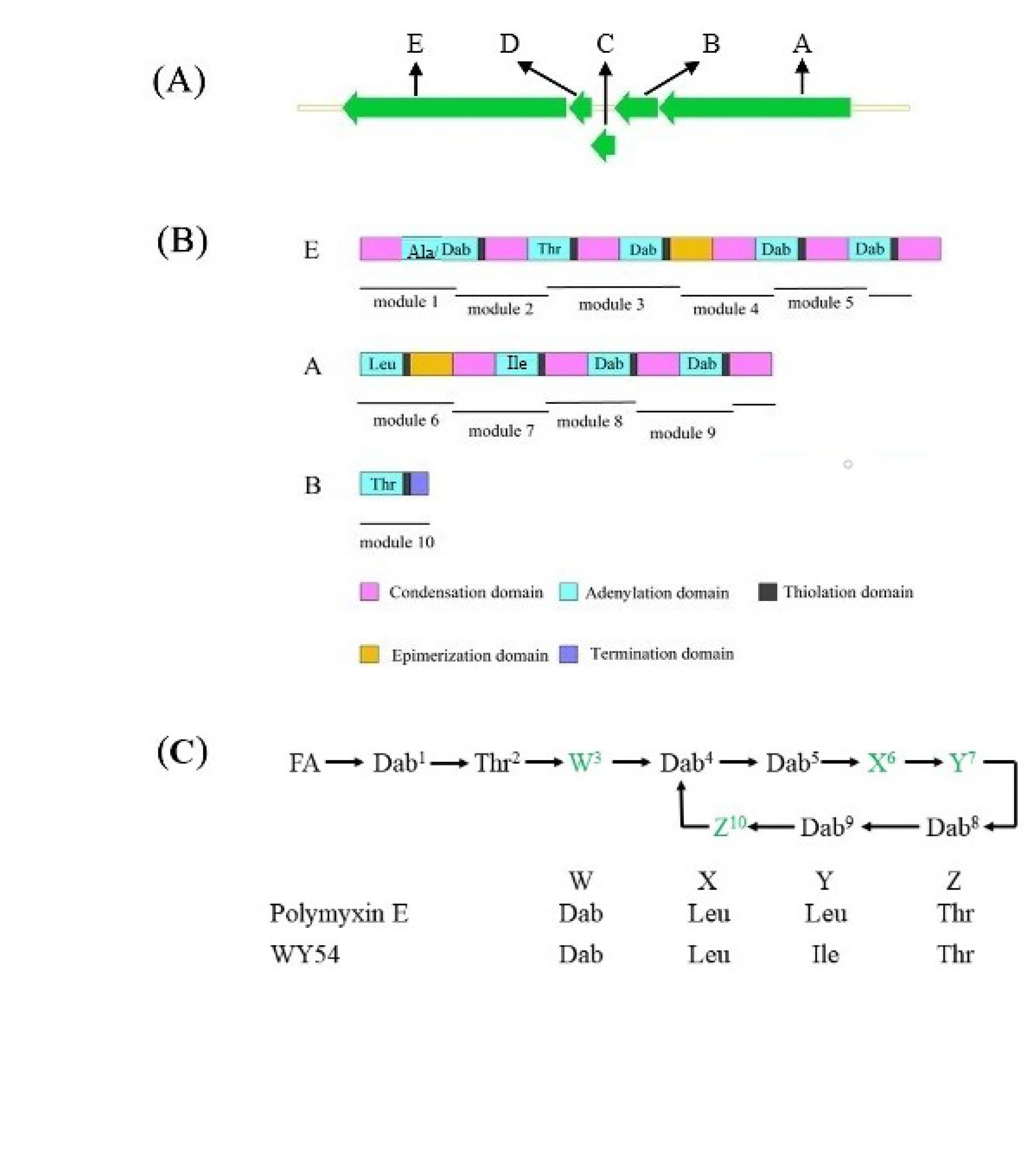
(A) The gene cluster responsible for the antagonistic compound; (B) The domain organization, and (C) The prediction of the core structure of the compound.

### Estimation of the molecular weight of the antagonistic compound

Concentrated samples of *P. polymyxa* WY54 wildtype strain and the WY54-MT7 mutant were separated by high performance liquid chromatography, respectively (Fig 6.A). Comparatively, the mutant strain WY54-MT7 lost a strong peak at the retention time of 9.437 min, which was eluted by 70% methanol to obtain a relatively high purity. Bioassay showed that the effluent from HPLC showed strong inhibition of *K. pneumonae* MH18(Fig 6. B), thus proving its antagonistic role. Then the concentrated sample of the antagonistic compound was further analyzed by mass spectrometry (Fig 6.C) and showed a molecular weight of 1168.38. This datum evidenced that the amino acid of the first position should be Dab, but not Ala, which was 29.04 smaller in molecular weight.

**Fig 6.**
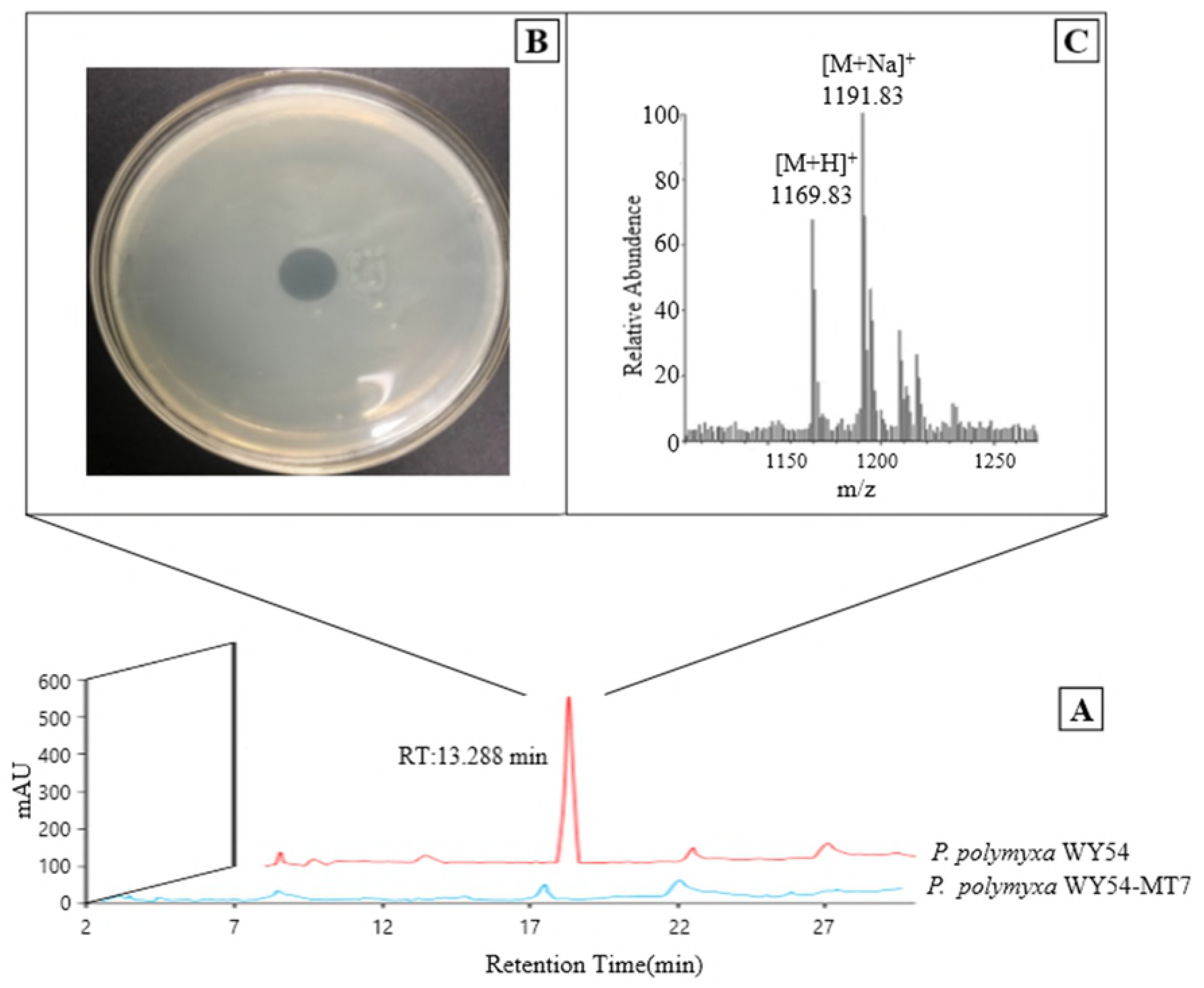
(A) Comparative HPLC analysis of the metabolites isolated from the wild-type strain WY54 and the mutant strain WY54-MT7. (B) Bioassay and (C) MS analysis of the peak that vanished in the null mutant from strain WY54.

## Discussion

Klebsiella pneumonia has been a major threat to animal industries and human health, the situation has been getting worse with the emergence of drug-resistant strains of *Klebsiella pneumoniae*(25–28), and requires novel strategies towards the prevention and treatment of this disease. By combining bacterial isolation and confrontation cultivation, we got a novel strain of *K. pneumoniae* that encodes the NDM-1 carbapenemase and was resistant to ceftriaxone sodium and meropenem. This strain *K. pneumoniae* MH18 caused an outbreak of pneumonia in a kennel of American bully, with 63.2% mortality. Meanwhile, a *Paenibacillus polymyxa* strain from the survival dogs was isolated to inhibit strain MH18. Drug-resistance coevolves with drug discovery, however, antibiotics remain the mainstay of anti-infection therapy(29, 30). Cephalosporins, fluoroquinolones and aminoglycosides were used as the primary weapons against *K. pneumoniae* infection(31), until extended-spectrum β-lactamases (ESBLs) that enabled *K. pneumoniae* to be resistant to various β-lactam antibiotics first found in 1983(32). Carbapenems were then developed to inhibit ESBL-producing pathogens for a decade, followed with the advent of carbapenem-resistant *K. pneumoniae* (CRKP) in 1993(33).

Fortunately, we isolated a strain of *Paenibacillus* from the feces of the healthy dogs in the kennel with the outbreak of Klebsiella infection, and the *Paenibacillus* strain demonstrated potent activity against the *K. pneumoniae* pathogen. *P. polymyxa* is well known for its ability to produce a variety of metabolites with antimicrobial activities(34–36). By combining transposon mutagenesis and bioinformatics analysis, we located the gene cluster involved in the antagonism. The gene cluster encoded a non-ribosomal peptide synthase, and antiSMASH data showed that the predicted structure of the product resembles that of polymyxin E1 with the first amino acid being Ala. Then we translated all the A-domains of the gene cluster for fine analysis with NRPSpredictor2, and revealed that the first adenylation domain recognized Dab. This was further proved by HPLC and MS analysis, with the molecular weight determined to be 1168.53. The fine structure and the mechanism of action of the compound remain to be discovered.

The gene clusters and their metabolites will allow us to decipher the gene expression and regulation towards genetic engineering and metabolic engineering. The data and procedure in this work opened a new avenue for the discovery and design of novel drugs against the superbugs.

## Acknowledgments

This work was in part supported by the National Natural Science Foundation of China (NSFC) Grant 31300111.

## Author Contributions

Conceived and designed the experiments: YW MY. Performed the experiments: MY HS. Analyzed the data: MY. Contributed reagents/materials/analysis tools: MY HL. Wrote the paper: MY XL XC.

